# SNAPPy: a snakemake pipeline for scalable HIV-1 subtyping by phylogenetic pairing

**DOI:** 10.1101/725838

**Authors:** Pedro M.M. Araújo, Joana S. Martins, Nuno S. Osório

**Author notes:** Corresponding author: Nuno S. Osório.

## Abstract

Human immunodeficiency virus 1 (HIV-1) genome sequencing is routinely done for drug resistance monitoring in hospitals worldwide. Subtyping these extensive datasets of HIV-1 sequences is a critical first step in molecular epidemiology and surveillance studies. The clinical relevance of HIV-1 subtypes is increasingly recognized. Several studies suggest subtype-related differences in disease progression, transmission route efficiency, immune evasion, and even therapeutic outcomes. HIV-1 subtyping is mainly done using web servers. These tools have limitations in scalability and potential noncompliance with data protection legislation. Thus, the aim of this work was to develop an efficient method for local and high-throughput HIV-1 subtyping. We designed SNAPPy: a snakemake pipeline for scalable HIV-1 subtyping by phylogenetic pairing. It contains several tasks of phylogenetic inference and BLAST queries, which can be executed sequentially or in parallel, taking advantage of multiple-core processing units. Although it was built for subtyping, SNAPPy is also useful to perform extensive HIV-1 alignments. This tool facilitates large-scale sequence-based HIV-1 research by providing a local, resource efficient and scalable alternative for HIV-1 subtyping. It is capable of analysing full-length genomes or partial HIV-1 genomic regions (GAG, POL, ENV) and recognizes more than 90 circulating recombinant forms. SNAPPy is freely available at: https://github.com/PMMAraujo/snappy.

## Introduction

The number of HIV-1 partial or complete genomic sequences in databases largely increased along the years after a noteworthy surge of almost tenfold in the 2000’s. HIV-1 genomic data is extremely valuable for fundamental research, translating into several epidemiological applications such as antiretroviral resistance surveillance or transmission history reconstruction [1–3]. Subtyping is a primary analysis done on HIV-1 sequences to allow further investigation.

HIV-1 was consensually divided in four groups (M, N, O and P), a consequence of multiple crossspecies transmission events from non-human primates to humans. M group is the only with a worldwide dispersion, and due to genetic differences and divergent evolutionary stories viruses from this group were divided into nine subtypes (A to D, F to H, J, and K) and sub-subtypes (e.g. A1, A2, F1, and F2). HIV-1 genomes composed of parts of different subtypes are known as recombinant forms, which can be named circulating recombinant forms (CRFs) if several cases are detected or unique recombinant forms (URFs) for more sporadic cases [4,5]. It has been reported that different HIV-1 subtypes may be better adapted for specific transmission routes [6,7], contain higher prevalence of polymorphisms known to influence immune systems [8,9] or antiretroviral treatment evasion [10–12], and lead to differences in the disease progression rate [2,13–15].

There are three main classes of approaches to perform HIV-1 subtyping: similarity-based (e.g. Stanford [16] and NCBI subtyping tool [17]); statistical-based (e.g. COMET [18], jpHMM [19], and STAR [20]); and phylogenetic-based (e.g. REGA [21] and SCUEAL [22]). Phylogenetic-based tools are considered the most sensitive and specific but also more time and computational resource consuming [21,23]. Most of the currently available tools are made available in the form of webservers, making them easy to access and use. Nevertheless, this distribution mode raises issues regarding the scalability of the implementation, it is unreasonable to provide a web-server without limitations in the input size or number of jobs. Making large scale analysis like multicentre molecular epidemiology studies, systematic reviews or databases curations practically impossible. Moreover, the HIV-1 genomic material corresponds to a clinical result and is often under data protection legislation as such, requiring in many cases an ethic approval for data sharing or submission in external servers.

Despite the large interest in using phylogeny for HIV-1 subtyping, existing tools have failed to address scalability and privacy issues. To answer these limitations, we used the Snakemake workflow management system [24] to create a reproducible and scalable HIV-1 subtyping method based on phylogenetic pairing (SNAPPy). By combining established tools with an innovative approach, this pipeline is capable of scaling according to the available computational resources, allowing the local analysis of large amounts (tens of thousands) of HIV-1 genomes. SNAPPy was built on top of the assumption that the phylogenetic relationship provides the best possible identification of the HIV-1 subtype [21,23]. However, recombination events represent exceptions to the assumption of a common ancestor (coalescent) [25]. Therefore, we complemented the phylogenetic inference with the similarity search method BLAST [26]. Reproducibility and efficient transmission of protocols are current challenges in bioinformatics, critical to share domain-specific knowledge [24,27]. Therefore, one of the main focus in SNAPPy is to give the user access to all the relevant intermediate files created and how the final subtyping decision was performed.

Overall, we present a problem-solving pipeline to allow local large-scale HIV-1 subtyping, based on phylogenetic inference and complemented with similarity search tasks.

## Implementation

### 1. SNAPPy architecture

The SNAPPy pipeline was built on the Snakemake workflow management system [24]. Several tools/software were used to perform different tasks within this pipeline: MAFFT v7.245 [28] for multiple sequence alignment; the Biopython v1.72 [29] modules SeqIO and Phylo [30] for data parsing and manipulation; BLAST v2.7.1 [26] for local database search; FastTree v2.1.10 [31] for phylogenetic inference. Other Python v3.6 [32] packages were also used to create tests, pytest [33], and data manipulation, numpy [33] and pandas [34]. For package management and to create a contained environment for SNAPPy, we recommend Conda [35], and provided a ready to use file to this end (‘environment.yaml’) as well as instructions on how to install and utilize it in SNAPPy’s documentation [36].

The term ‘target’ used in this manuscript refers to the file currently being processed by SNAPPy. When used for subtyping, SNAPPy performs the following tasks to a given target sequence: 1) split the input in multiple single FASTA files; 2) alignment to the reference genome; 3) BLAST against a set of HIV-1 reference sequences; 4) perform phylogenetic inferences using the BLAST top hits, the target, and an outgroup sequence; 5) sliding window BLAST against a database of HIV-1 reference sequences; 6) concatenation and analysis of the results obtained in the previous tasks and creation of the output results. The Figure 1 is a schematic representation of this pipeline. At the end of the subtyping task, SNAPPy produces two files the ‘subtype_results.csv’ and the ‘report_subtype_results.csv’, corresponding to a simplified version of the subtyping result and a more extensive report of all the outputs created by SNAPPy.

Alternatively, as any other Snakemake [24] pipeline, intermediate tasks can be performed without the execution of the entire pipeline, making SNAPPy extremely useful for HIV-1 multiple sequence alignment (Figure 1, point 7). To match the names of the intermediate files created by SNAPPy and the header of the target sequences, a file named ‘keys_and_ids.csv’ is created. An in-depth description of each of these general steps can be seen in the following topics.

**Figure 1:**
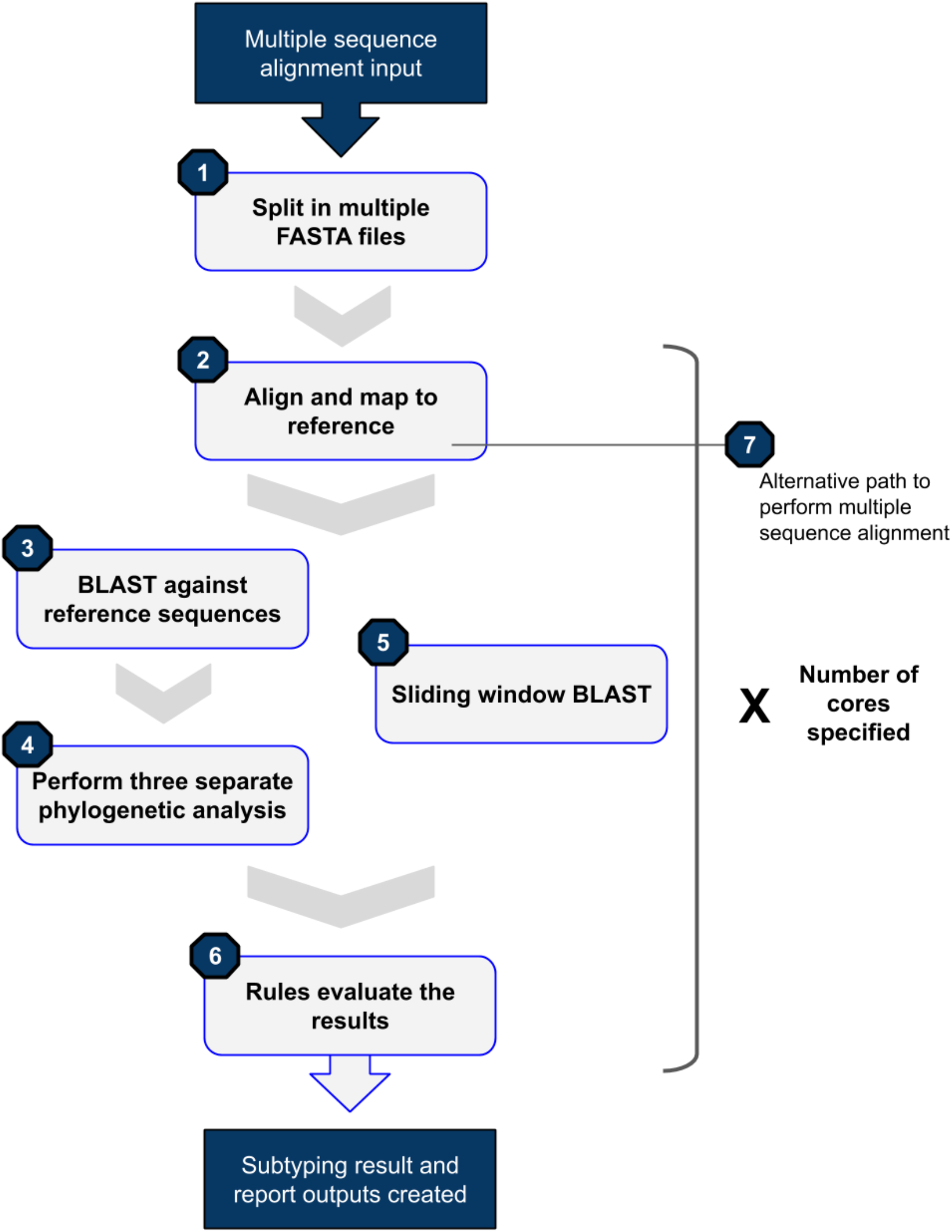
SNAPPy workflow diagram

#### 1.1. Reference Sequences

In all instances of SNAPPy, the HXB2 (GenBank: K03455) reference genome was used as a genomic position reference. The outgroup sequence used in the phylogenetic analysis corresponds to the CONSENSUS_CPZ sequence from the HIV sequence database [37] (Alignment type: Consensus/Ancestral, Year: 2002, Organism: Other SIV, Alignment ID: S02CG1). The creation of a comprehensive set of HIV-1 reference sequences proved to be a challenge. We based our dataset creation on the previously curated ‘HIV sequence database 2010 subtype reference genomes sequences compendium’ [38] and the ‘HIV Drug Resistance Database reference sequences’ [39]. The final subtype reference dataset for SNAPPy consisted of 493 genomic sequences. It included references for groups N, O, P and within group M for subtypes B, C, D, G, H, J and K, sub-subtypes A1, A2, F1 and F2 and CRFs until the number 96. Please notice that not all CRFs are represented due to lack of reference sequence or quality in the available data. The full reference dataset (493 sequences) is used for the BLAST task (see topic 1.3.) and a subset only containing groups, subtypes, sub-subtypes and CRFs 01 and 02 references (56 sequences) is used in the sliding window BLAST (see topic 1.5. Sliding window BLAST). For more information on these reference datasets please consult the Table 1 of Supplementary data.

#### 1.2. Alignment to reference

After splitting the multiple sequence alignment (MSA) into several single sequence FASTA files, each of them is aligned to the HIV-1 reference genome (HXB2). The module SeqIO from Biopython [29] is used to parse and manipulate the FASTA files in SNAPPy. The alignment is done using MAFFT [28]. The alignment method used does not allow the insertion of gaps in the reference sequence. After the alignment is performed, the target sequence is trimmed to only contain the genomic region specified by the user in the ‘config.yaml’ file. Being the currently available options ‘GAG’, ‘PR’, ‘RT’, ‘PR-RT’, ‘INT’, ‘POL’, ‘ENV’ and ‘GAG-POL-ENV’, which correspond to the HIV-1 genomic regions with the same names in the HXB2 reference genome. The resulting files are them written to the ‘aligned’ folder.

#### 1.3. BLAST

The obtained alignments are them BLASTed against a local database of 493 HIV-1 reference sequences (see topic 1.1.). For this task, BLAST [26] is used. The results were sorted by bitscore (considering higher is better) and the best scoring result is outputted in the ‘report_subtype_results.csv’ file in the column ‘closser_ref’. The BLAST results are also used to make three groups of references sequences: containing the first 48 results; containing the first 48 results of only subtype references; containing the first 48 results of only CRF references. These three groups of reference sequences are then used in the phylogenetic analysis. The selected number of sequences (48) showed good compromise between analysis time and reproducibility, as discussed in topic 1.4. Since this BLAST task is done as preparation step to the phylogenetic analysis, it must have high sensitivity, so no related reference sequences are missed. To achieve this, the word size parameter was set to 10. We also restricted the cutoff E-value to 1.0e-10 to avoid the creation of large output files, without restricting the results. The intermediate files of the BLAST analysis are outputted to the ‘blast’ folder, being available for further consulting. For the split in subtype and CRF references in this step of SNAPPy, CRFs 01 and 02 are treated as subtypes, not CRFs. This decision was made based on the high prevalence of these CRFs and their ambiguous origin [40,41].

#### 1.4. Phylogenetic inference

To the three previously selected groups of 48 references (see topic 1.2.), the target sequence and a non-HIV-1 sequence for rooting (see topic 1.1.) are added. Obtaining three sets of 50 sequences that will serve as inputs for the phylogenetic analysis. Groups of 50 sequences showed to be contained and yet a comprehensive set of sequences to perform the phylogenetic inference. To perform the phylogenetic analysis, FastTree [31] was used with the GTR nucleotide substitution model. The Biopython module Phylo [30] was applied to parse and manipulate the phylogenetic trees created within SNAPPy. After rooting on the outgroup, it is inferred if the target sequence belongs to a monophyletic clade with sequences of one, and only one, subtype or CRF. If this happened, we consider that there is phylogenetic evidence of the relationship between the target sequence and a reference subtype/CRF. The result of this inference is, together with the support values for that node (Shimodaira-Hasegawa test, as implemented in FastTree), are then outputted to the ‘report_subtype_results.csv’ file. Resulting in six output columns: ‘node_all_refs’, ‘s_node_all_refs’, ‘node_pure_refs’, ‘s_node_pure_refs’, ‘node_recomb_refs’ and ‘s_node_recomb_refs’. The intermediate files of the phylogenetic analysis are outputted to the ‘trees’ folder. The notation ‘all’, ‘pure’, and ‘recomb’ refers to the set of references used for that phylogenetic reconstruction.

#### 1.5. Sliding window BLAST

The sliding window BLAST can be performed in parallel with the tasks described in topics 1.3. and 1.4., being only dependent on the outputs from the Alignment to the reference task (topic 1.2.). Initially, the positions in the target sequence corresponding to gaps (‘-’) are excluded. For this task, BLAST [26] is used. The length of the sliding window used is 400 nucleotides, and smaller fragments/sequences are not processed by this method. The step size used is 50 nucleotides, creating eight bins for each window. The result for each BLAST window, and consequently its eight bins, is the subtype of the top result (bitscore) reference sequence. If more than one sequence of different subtypes has the same top score, the output for all bins of that window is null (‘-’). If the method fails to produce an output, the result for all bins of that window is null. After all possible sliding windows have been BLASTed, several bins will have multiple outputs. Then, a majority rule is applied to decide the final subtype for that bin. In case of a tie, the result for that bin is null. Figure 2 contains a schematic representation of this process. The database used to BLAST against in this task only contains the group, subtype, sub-subtype and CRF1 01 and 02 references, as described in the references section (56 sequences). The word size parameter applied was 30, with the purpose of obtaining high specificity. Values higher than this showed to cause instability in our tests (low reproducibility). An E-value cutoff of 1.e-50 was used to ensure the generated outputs were not too large, without losing real BLAST hits. The outputs of this inference are written to the ‘report_subtype_results.csv’ file in the column ‘recomb_result’ and the resulting files from these tasks are outputted to the ‘blast’ folder.

**Figure 2:**
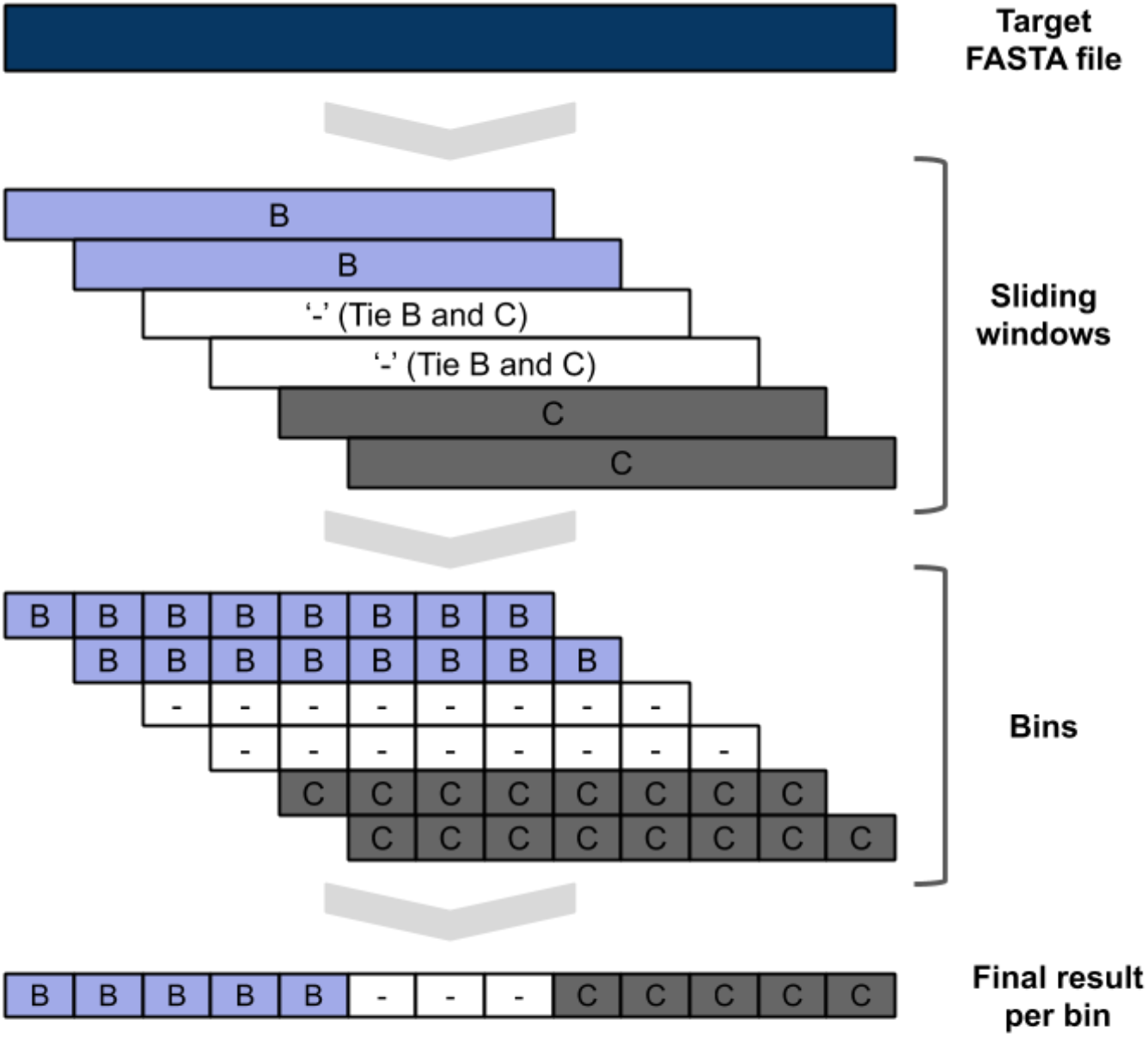
Sliding window BLAST schematic representation

#### 1.6. Decision Rules

The results generated by the full sequence BLAST, phylogenetic analysis, and the sliding window BLAST are then processed using a set of rules to produce the final output. These rules are executed in order, meaning that the second rule is only applied if the first rule criteria were not met, and so forth. The list of these rules can be consulted in Table 2 of Supplementary data. At the end of the process, two final outputs are created in the snappy folder: ‘subtype_results.csv’ and ‘report_subtype_results.csv’, as mentioned above.

#### 1.7. System management

SNAPPy is a pipeline that performs several tasks, some of them generate a relatively large number of outputs. Therefore, in order to avoid wasting unnecessary disk space and simplifying the user experience, some of the intermediate files produced by SNAPPy are deleted before the end of the process. However, all the relevant files for the subtyping decision are kept and available for consulting after the pipeline finishes. At the end of each SNAPPy run, a snakemake hidden folder named ‘.snakemake’ is deleted because it occupies a substantial amount of space. However, this folder contains all the logs about the tasks performed and may be useful for debugging.

SNAPPy is distributed with a series of built-in tests created using pytest [33]. After installing SNAPPy or after making alterations in SNAPPy’s folder is recommended to run the tests to infer if the pipeline is behaving as expected.

SNAPPy’s documentation [36] includes detailed instructions on: pipeline installation; tutorials on how to use it; a list of available commands; an in-depth description of each pipeline step; FAQ; and how to cite and user license information.

### 2. Pipeline evaluation

#### 2.1. Reproducibility

One of the bases of SNAPPy is phylogenetic inference, which has some stochasticity involved. As implemented in FastTree [31], the initial topology is constructed based on heuristic methods, which afterwards is optimized with maximum-likelihood rearrangements. In theory, this could lead to variance in the output of the subtyping pipeline. Furthermore, for the evaluation of the branch support, we were faced with the possibility of using statistic-based (e.g. Shimodaira-Hasegawa test) or sampling-based (e.g. bootstrapping) methods. In our tests, we found that the usage of bootstrapping approaches leads to some lack of reproducibility in the pipeline outputs, even at one thousand replicates, as previously reported [21]. Which may be explained by the low percentage of informative sites found in the tested HIV-1 sequences, which are extremely similar among each other. The increment of the bootstrapping replicates would lead to an exponential increment of the phylogenetic inference step computational time, making the pipeline much slower. Therefore, we decided to use a statistic-based branch support inference method, the Shimodaira-Hasegawa test, for phylogenetic inference in SNAPPy. As stated in the topics 1.3. and 1.5., when using BLAST, the word size and cutoff parameters were selected to achieve the desired objectives (sensitivity or specificity) and ensure the stability of the analysis. After the pipeline was constructed, we performed 6 sets of 3 independent SNAPPy runs with a test set of 5285 sequences (see topic 2.2.) and compared the outputs of each independent run in terms of reproducibility. The obtained result was 100% reproducibility, meaning that the output file ‘subtyping_results.csv’ for each of the independent runs were exactly the same.

#### 2.2. Scalability

SNAPPy was built for large-scale analysis, taking advantage of modern multi core/thread CPUs. The usage of Snakemake [24] as a workflow manager allows the construction of a directed acyclic graph (DAG) of jobs, inferring which tasks need to be performed sequential and which can run in parallel. To infer the overall scalability of SNAPPy regarding multithreading, we performed the subtyping of a test set of 5285 sequences with the following number of cpu threads: 1; 2; 4; 8; 16; and 32. The selection of these numbers was made having as objective the comparison of the computation time reduction by half (halving) when doubling the amount of computational resources. In Figure 3, there is a comparison of the real time that SNAPPy took to subtype the test set versus the expected halving time. This expected time reduction is purely theoretical and constructed based on the time SNAPPy took to subtype the test set with one core and subsequence duplication of the number of computational resources used. These tests were performed in a server with double Xeon E5-2680 2.50 GHz cpu (12 cores/24 threads), 128 GB of ram 2133 MHz, in a SATA III SSD hard drive.

**Figure 3:**
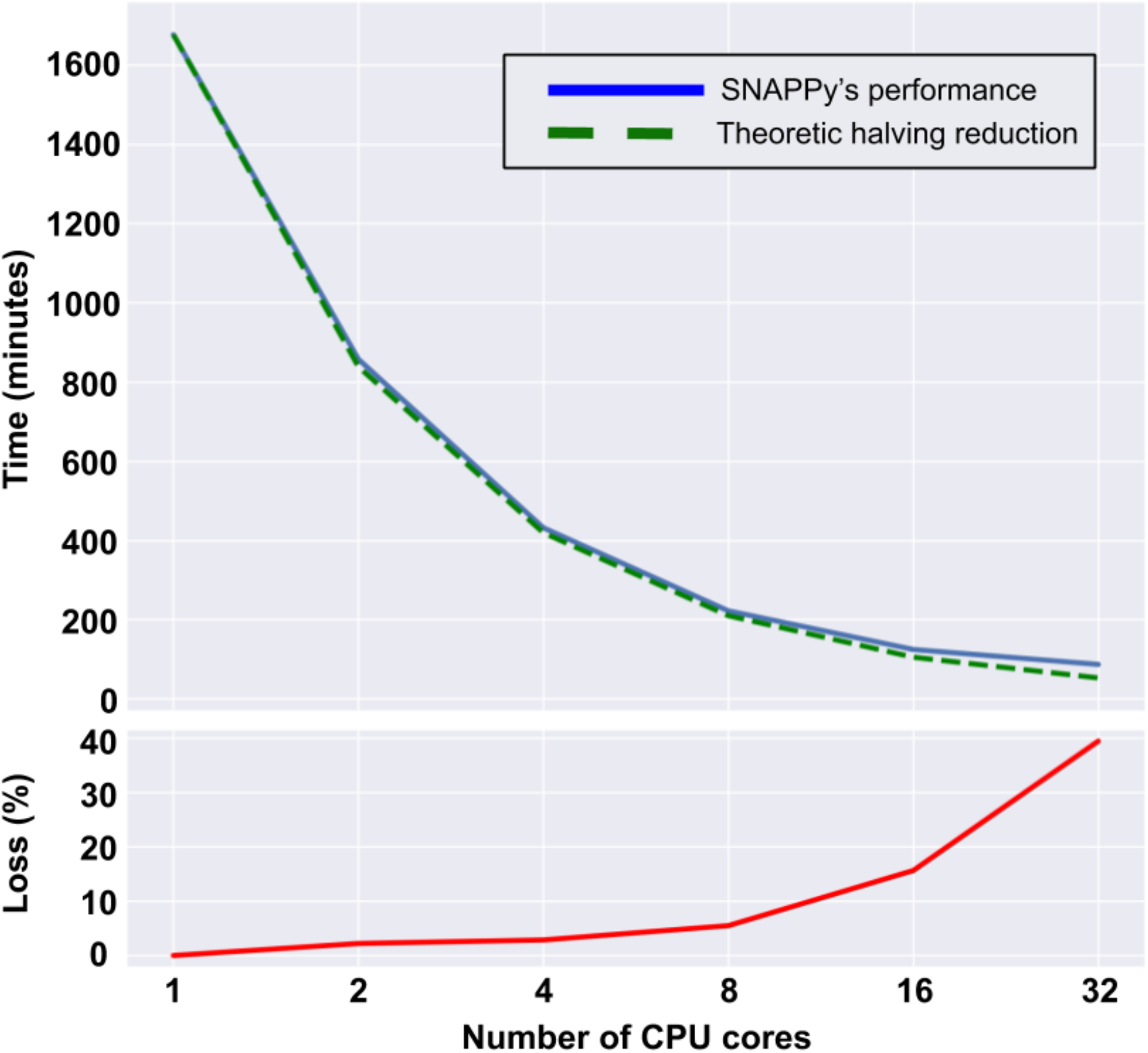
Multiplethread CPU time performance and associated percentage loss

SNAPPy manages the generated files in order to give the user all the information needed to understand the results and decisions made. This feature is a trade-off purposely made to give users the maximum amount of information without wasting disk space. Nevertheless, when used for large scale analysis (tens of thousands of sequences), SNAPPy will create a large number of small files that together occupy a considerable amount of disk space. As an indicator, a SNAPPy run of 50 thousand HIV-1 sequences occupied at the peak 59 GB and less than 4 GB after the depletion of the snakemake hidden logs folder.

#### 2.3. Subtyping methods comparison

The division of HIV-1 in groups, subtypes, and sub-subtypes is extremely valuable for epidemiological inferences [1–3]. However, this division is a man-made construction that only makes sense in the eyes of a phylogenetic or epidemy reconstruction [2,5]. Therefore, it makes sense to argue that phylogenetic reconstruction is the gold standard for HIV-1 subtyping [21,23], with the exception of recombination events that represent a deviation from the coalescent assumption [25]. Since SNAPPy is based on phylogenetic inference, using phylogeny to evaluate SNAPPy’s performance would be poorly informative. Given this limitation, we decided to test SNAPPy against a set of other HIV-1 subtyping methods, evaluating their convergence and divergence. The selected HIV-1 subtyping methods were REGA v3.0 [21], COMET v2.3 [18] and SCUEAL [22]. This selection was made based on our best knowledge of available tools including statistical and phylogeny-based methods.

For this comparison, a test set of 5285 sequences was created. From all the available complete HIV-1 genomes (>9100) in the HIV-1 sequence database [37] 10% of sequences for each of the subtypes present (according to the database) were selected at random, comprising a total of 1057 genomes. Those genomes were then trimmed for 4 genomic regions: ENV; GAG; POL; and PR-RT. This test set was designed to explore the capabilities of each subtyping method in an extensive variety of subtypes while using different HIV-1 genomic regions as input. However, there are differences in the methods implementations; SCUEAL only subtypes sequences of the POL region, therefore the comparison dataset for this tool is smaller (1057 sequences x 3 regions = 3171 sequences), and is capable of recognizing CRFs until the number 43; REGA is able to recognize CRFs until the number 47; COMET is cable of recognizing CRFs until the number 96. The outputs for each subtyping tool required some manipulation in order to achieve a “common language”, making it possible to compare all the tools outputs. Regarding REGA outputs, the terms “like” and “potential recombinant” were excluded, maintaining the assigned subtype; the outputs with “recombination of” were named URFs of the described subtypes or URF_CPX if there was an indication of more than two recombining subtypes. REGA results marked only with the information of “Recombinant” were transformed into URF_CPX; and when the outputs was “Check the report” no subtyping result was assigned. For the COMET outputs the only transformation was to change “unassigned” to URFs of the subtypes present or URF_CPX if more than two subtypes were reported. Concerning the SCUEAL outputs, the words “ancestral” and “like” were excluded; “Complex” was converted to URF_CPX and “AE” to CRF_01. The SCUEL results with “recombinant” and more than two subtypes were converted to URF_CPX and those with less than 2 subtypes converted to URFs; for outputs with “U” and “FAILED” no subtyping result was assigned. The comparison between the four subtyping methods results can be seen in Figure 4. The highest level of agreement was observed between SNAPPy and COMET (83%), while SCUEAL and REGA had the lowest concordance (61%). The remaining pairs showed results in the range between 72% and 78%.

**Figure 4:**
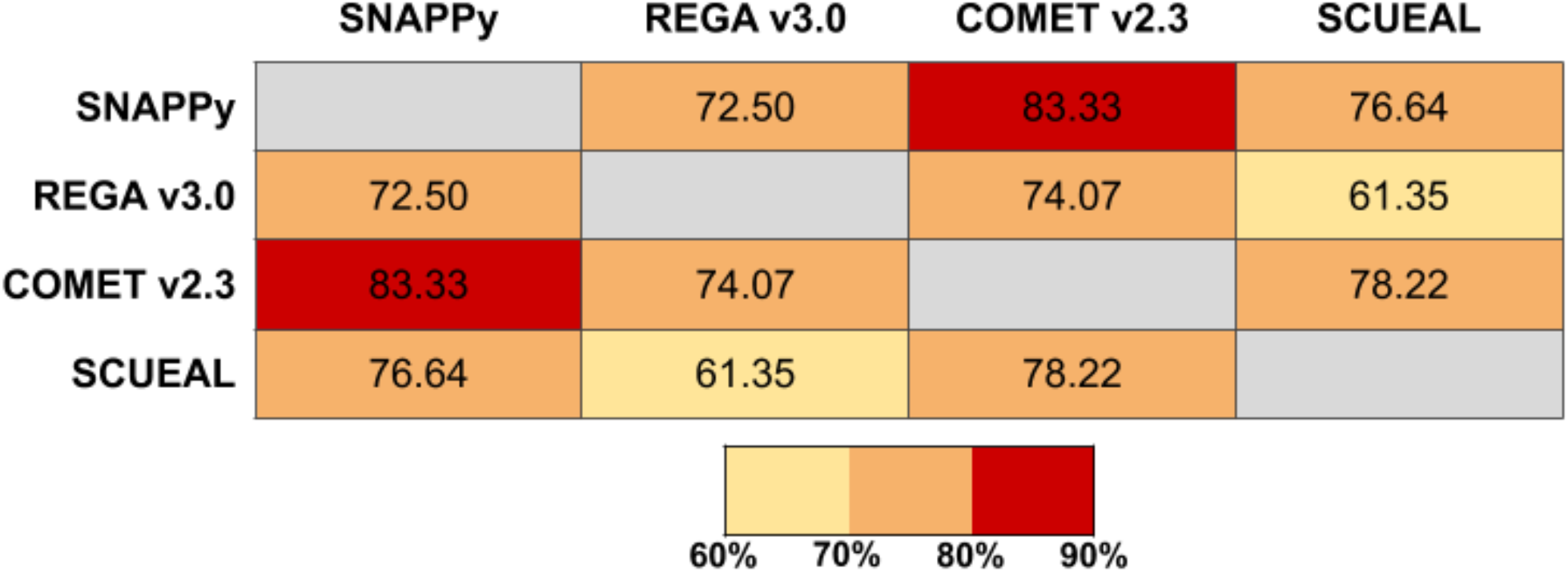
Percentage of agreement observed among the HIV-1 subtyping tools tested

We also calculated the precision, recall, and F1 scores (balance of precision and recall) for the three subtyping methods tested (REGA, SCUEAL and COMET) versus SNAPPy (Supplementary Table 3 to 5). This analysis was performed to give an indication for users comparing results obtained with other tools and SNAPPy, for each HIV-1 group, subtype, sub-subtype, CRF, and URF. The precision and recall metrics can be seen as indicators if SNAPPy is classifying a given subtype in the test set more or less often, respectively, than the subtyping method it is being compared with. Without surprise, in the test set the results for subtypes B and C (the most abundant) and non-M HIV-1 groups (N, O and P) showed the highest F1 scores. The results for URFs and CRFs showed great variability.

## Discussion and Conclusions

The quantity of available HIV-1 genomes is ever increasing, the manual handling of such large amounts of data is impractical, leading to the need of creating analysis pipelines. Such pipelines targeting specific challenges are a practical and effective way of disseminating domain knowledge and increasing reproducibility [24,27]. The test set used for the different metrics evaluated here is composed of 1057 sequences of HIV-1 genomes and the same sequences trimmed for the genomic regions ENV, GAG, POL, and PR-RT corresponding to a total of 5285 sequences.

There is some stochasticity involved in the phylogenetic inference process as established in FastTree [31]. Nevertheless, the parameters selected for the branching support evaluation and the number of samples per tree allowed 100% reproducibility among independent SNAPPy runs results in an extremely diverse test set. This outcome highlights the versatility and reliability of this pipeline.

Together with the increment in the amount of data available, there have been hardware improvements, particularly in recent years CPUs with a high number of cores/threads (≧8) have reached the mainstream segment of the market. Therefore, tool building should be done to take maximum advantage of these resources. Snakemake [24], the pipeline workflow management systems that SNAPPy is built upon, allows an almost linear scaling in the ratio of computational time/ number of CPU cores used. The expected reduction by half of the computational time by doubling the number of threads used was observed in SNAPPy runs of the test set with minor percentage lost when using 2, 4, 8, or 16 CPU threads (2%, 3%, 5%, and 16% respectively). However, for the runs with 32 threads, the drop from the expected halving time was almost 40%. This drop may be a consequence of the hardware used, since it is composed of 24 independent CPU cores (2 cpus x 12 cores) passing that number may cause single thread performance loss. Therefore, we do not advice the usage of SNAPPy multithreading capabilities in more instances than the number of physical CPU cores of the machine used.

The classification of different HIV-1 sequences in groups, subtypes, sub-subtypes or recombinant forms is a challenging and sometimes ambiguous process. Sequence-based phylogenetic reconstruction of the evolutionary history assuming a common ancestry of the viral samples is consensually identified as the best approach for HIV-1 subtyping [21,23]. However, the coalescent assumption is not fulfilled in the cases of recombination [25]. Therefore, two complementary approaches were used in SNAPPY, one based on BLAST [26] similarity searches and another based on the phylogenetic inference (FastTree [31]).

The side-by-side comparison of different HIV-1 subtyping methodologies is complex and sometimes impossible. Here, we compared SNAPPy, REGA [21], COMET [18], and SCUEAL [22] outputs for a test set of 5285 sequences (Figure 3). As described in the methods section, the regions of the HIV-1 genome these tools are capable of subtyping and the number of CRFs they are able to identify varies. Without surprise, COMET and SNAPPy have the highest value of accordance (>80%) among the tested tools. This outcome is highly influenced by the fact that these tools are prepared to identify a large number of CRFs (> 90) in comparison with the remaining two tools in this test. The lowest accordance was observed between the pair REGA and SCUEAL (61%). The lowest pairing for SNAPPy was REGA with 73% followed by SCUEAL with 77%.

In Supplementary tables 3 to 5, we show the precision, recall, and F1 scores resulting from the comparison of three subtyping methods (REGA, COMET and SCUEAL) with SNAPPy, for each HIV-1 group, subtype, sub-subtype, CRF, and URF. The overall F1 scores for REGA and SCUEAL suffer from the fact that these tools identify a narrower range of CRFs than SNAPPy. Moreover, SCUEAL only subtypes sequences for the POL HIV-1 genomic region, being sequences from the remaining regions treated as missing data, and therefore driving the F1 score further down. Nevertheless, is it expected a great reproducibility among these two tools and SNAPPy for subtypes B and C, group N and several CRFs (01, 05, 13, 27, 35, 40, 42 and 47 for REGA and 05, 17, 18, 19, 24, 31, and 33 for SCUEAL). On the other hand, COMET is capable of identifying a wide range of CRFs, similarly to SNAPPy, which is observed in the overall F1 score (0.83), precision (0.88), and recall (0.83) results. Regarding the test set, these two tools identified CRFs 17, 27, 34, 40, 47, 68, and 74 in exactly the same cases and subtypes B and C, and groups N, P and O with a high similarity rate (F1 scores respectively: 0.99, 0.94, 1.0, 1.0, and 0.96). The results for subtypes that showed less reproducibility among the three tested methods and SNAPPy, were subtypes H and K and sub-subtype A2 and F2.

These results highlight the variability observed among HIV-1 subtyping tools, which is expected [21,23,42] and should not be seen as a drawback but instead as an interval of possibilities around a subtyping result. Moreover, these results demonstrate that when several HIV-1 tools agree in one result, there is a high degree of confidence in that outcome. SNAPPy is not strictly better or worse than the other tools regarding the final result, but it is a needed addition to this space, allowing local high-throughput HIV-1 subtyping while being versatile, reliable and cable of scaling. The results reported here emphasized the importance of SNAPPy to facilitate the subtype annotation of large datasets of HIV-1 genomic sequences. This work represents a novel approach for HIV-1 subtyping that can contribute significantly towards a better understanding of the relevant roles and traits of the different HIV-1 subtypes.

Key points:

- The amount of available HIV-1 genomic information is increasing, therefore there is a need to create tools to perform large-scale analysis of HIV-1 subtypes.
- HIV-1 subtyping methods have considerable differences in their implementations and, consequently, their outputs.
- SNAPPy is capable of local high-throughput HIV-1 subtyping with great reproducibility while being able to scale according to the computational resources available.

## Author Contributions

PMMA and NSO conceived the pipeline. The pipeline and documentation were constructed by PMMA and critically assessed by JSM. The manuscript was written by PMMA and reviewed by JSM and NSO.

## Availability

SNAPPy source code is freely available via GitHub at: https://github.com/PMMAraujo/snappy. SNAPPy documentation can be consulted at: https://snappy-hiv1-subtyping.readthedocs.io/.

## Supplementary Data

(in separated document)

## References

1. Yebra G, Ragonnet-Cronin M, Ssemwanga D, et al. Analysis of the history and spread of HIV-1 in Uganda using phylodynamics. J. Gen. Virol. 2015; 96:1890–8

2. Araujo PMM, Carvalho A, Pingarilho M, et al. Characterization of a large cluster of HIV-1 A1 infections detected in Portugal and connected to several Western European countries. Sci. Rep. 2019; 9:7223

3. Abecasis AB, Wensing AMJ, Paraskevis D, et al. HIV-1 subtype distribution and its demographic determinants in newly diagnosed patients in Europe suggest highly compartmentalized epidemics. Retrovirology 2013; 10:7

4. Robertson DL, Anderson JP, Bradac JA, et al. HIV-1 nomenclature proposal. Science 2000; 288:55–6

5. Hemelaar J. Implications of HIV diversity for the HIV-1 pandemic. J. Infect. 2013; 66:391–400

6. Renjifo B, Gilbert P, Chaplin B, et al. Preferential in-utero transmission of HIV-1 subtype C as compared to HIV-1 subtype A or D. AIDS 2004; 18:1629–36

7. John-Stewart GC, Nduati RW, Rousseau CM, et al. Subtype C Is associated with increased vaginal shedding of HIV-1. J. Infect. Dis. 2005; 192:492–6

8. Serwanga J, Nakiboneka R, Mugaba S, et al. Frequencies of Gag-restricted T-cell escape ‘footprints’ differ across HIV-1 clades A1 and D chronically infected Ugandans irrespective of host HLA B alleles. Vaccine 2015; 33:1664–72

9. Bartolo I, Abecasis AB, Borrego P, et al. Origin and epidemiological history of HIV-1 CRF14_BG. PLoS One 2011; 6:e24130

10. Camacho RJ, Vandamme A-M. Antiretroviral resistance in different HIV-1 subtypes: impact on therapy outcomes and resistance testing interpretation. Curr. Opin. HIV AIDS 2007; 2:123–9

11. Abecasis AB, Deforche K, Snoeck J, et al. Protease mutation M89I/V is linked to therapy failure in patients infected with the HIV-1 non-B subtypes C, F or G. AIDS 2005; 19:1799–806

12. Brenner B, Turner D, Oliveira M, et al. A V106M mutation in HIV-1 clade C viruses exposed to efavirenz confers cross-resistance to non-nucleoside reverse transcriptase inhibitors. AIDS 2003; 17:F1-5

13. Kiwanuka N, Laeyendecker O, Robb M, et al. Effect of human immunodeficiency virus Type 1 (HIV-1) subtype on disease progression in persons from Rakai, Uganda, with incident HIV-1 infection. J. Infect. Dis. 2008; 197:707–13

14. Easterbrook PJ, Smith M, Mullen J, et al. Impact of HIV-1 viral subtype on disease progression and response to antiretroviral therapy. J. Int. AIDS Soc. 2010; 13:4

15. Baeten JM, Chohan B, Lavreys L, et al. HIV-1 subtype D infection is associated with faster disease progression than subtype A in spite of similar plasma HIV-1 loads. J. Infect. Dis. 2007; 195:1177–80

16. Liu TF, Shafer RW. Web resources for HIV type 1 genotypic-resistance test interpretation. Clin. Infect. Dis. 2006; 42:1608–18

17. Rozanov M, Plikat U, Chappey C, et al. A web-based genotyping resource for viral sequences. Nucleic Acids Res. 2004; 32:W654-9

18. Struck D, Lawyer G, Ternes AM, et al. COMET: Adaptive context-based modeling for ultrafast HIV-1 subtype identification. Nucleic Acids Res. 2014; 42:e144

19. Schultz A-K, Zhang M, Bulla I, et al. jpHMM: Improving the reliability of recombination prediction in HIV-1. Nucleic Acids Res. 2009; 37:W647–51

20. Myers RE, Gale C V, Harrison A, et al. A statistical model for HIV-1 sequence classification using the subtype analyser (STAR). Bioinformatics 2005; 21:3535–40

21. Pineda-Pena A-C, Faria NR, Imbrechts S, et al. Automated subtyping of HIV-1 genetic sequences for clinical and surveillance purposes: performance evaluation of the new REGA version 3 and seven other tools. Infect. Genet. Evol. 2013; 19:337–48

22. Kosakovsky Pond SL, Posada D, Stawiski E, et al. An evolutionary model-based algorithm for accurate phylogenetic breakpoint mapping and subtype prediction in HIV-1. PLoS Comput. Biol. 2009; 5:e1000581

23. Fabeni L, Berno G, Fokam J, et al. Comparative Evaluation of Subtyping Tools for Surveillance of Newly Emerging HIV-1 Strains. J. Clin. Microbiol. 2017; 55:2827–2837

24. Koster J, Rahmann S. Snakemake--a scalable bioinformatics workflow engine. Bioinformatics 2012; 28:2520–2

25. Pérez-Losada M, Arenas M, Galán JC, et al. Recombination in viruses: Mechanisms, methods of study, and evolutionary consequences. Infect. Genet. Evol. 2015; 30:296–307

26. Camacho C, Coulouris G, Avagyan V, et al. BLAST+: architecture and applications. BMC Bioinformatics 2009; 10:421

27. Di Tommaso P, Chatzou M, Floden EW, et al. Nextflow enables reproducible computational workflows. Nat. Biotechnol. 2017; 35:316–319

28. Katoh K, Standley DM. MAFFT multiple sequence alignment software version 7: improvements in performance and usability. Mol. Biol. Evol. 2013; 30:772–80

29. Cock PJA, Antao T, Chang JT, et al. Biopython: freely available Python tools for computational molecular biology and bioinformatics. Bioinformatics 2009; 25:1422–3

30. Talevich E, Invergo BM, Cock PJA, et al. Bio.Phylo: a unified toolkit for processing, analyzing and visualizing phylogenetic trees in Biopython. BMC Bioinformatics 2012; 13:209

31. Price MN, Dehal PS, Arkin AP. FastTree 2--approximately maximum-likelihood trees for large alignments. PLoS One 2010; 5:e9490

32. Van Rossum G, Drake FL. Python 3 Reference Manual. 2009

33. Pytest. Computer software. Vers. 4.5.0. https://docs.pytest.org/en/latest/

34. McKinney W. Data Structures for Statistical Computing in Python. Proc. 9th Python Sci. Conf. 2010; 51–56

35. Anaconda Software Distribution. Computer software. Vers. 3-4.6.14. Miniconda, Apr. 2019. https://anaconda.com

36. Araújo PMM. SNAPPy’s documentation: https://snappy-hiv1-subtyping.readthedocs.io/en/latest/

37. Theoretical Biology and Biophysics Group, Los Alamos National Laboratory, HIV sequence database. http://www.hiv.lanl.gov/

38. Kuiken C, Foley B, Leitner T, et al. HIV Sequence Compendium 2010. Eds. Published by Theoretical Biology and Biophysics Group, Los Alamos National Laboratory, NM, LA-UR 10–03684

39. Shafer RW. Rationale and uses of a public HIV drug-resistance database. J. Infect. Dis. 2006; 194 Suppl:S51–8

40. Gao F, Robertson DL, Morrison SG, et al. The heterosexual human immunodeficiency virus type 1 epidemic in Thailand is caused by an intersubtype (A/E) recombinant of African origin. J. Virol. 1996; 70:7013–29

41. Abecasis AB, Lemey P, Vidal N, et al. Recombination confounds the early evolutionary history of human immunodeficiency virus type 1: subtype G is a circulating recombinant form. J. Virol. 2007; 81:8543–51

42. Gifford R, de Oliveira T, Rambaut A, et al. Assessment of automated genotyping protocols as tools for surveillance of HIV-1 genetic diversity. AIDS 2006; 20:1521–9

